# Analysis of functional connectivity and oscillatory power using DICS: from raw MEG data to group-level statistics in Python

**DOI:** 10.1101/245530

**Authors:** Marijn van Vliet, Mia Liljeström, Susanna Aro, Riitta Salmelin, Jan Kujala

## Abstract

Communication between brain regions is thought to be facilitated by the synchronization of oscillatory activity. Hence, large-scale functional networks within the brain may be estimated by measuring synchronicity between regions. Neurophysiological recordings, such as magnetoencephalography (MEG) and electroencephalography (EEG), provide a direct measure of oscillatory neural activity with millisecond temporal resolution. In this paper, we describe a full data analysis pipeline for functional connectivity analysis based on dynamic imaging of coherent sources (DICS) of MEG data. DICS is a beamforming technique in the frequency-domain that enables the study of the cortical sources of oscillatory activity and synchronization between brain regions. All the analysis steps, starting from the raw MEG data up to publication-ready group-level statistics and visualization, are discussed in depth, including methodological considerations, rules of thumb and tradeoffs. We start by computing cross-spectral density (CSD) matrices using a wavelet approach in several frequency bands (alpha, theta, beta, gamma). We then provide a way to create comparable source spaces across subjects and discuss the cortical mapping of spectral power. For connectivity analysis, we present a canonical computation of coherence that facilitates a stable estimation of all-to-all connectivity. Finally, we use group-level statistics to limit the network to cortical regions for which significant differences between experimental conditions are detected and produce vertex-and parcel-level visualizations of the different brain networks. Code examples using the MNE-Python package are provided at each step, guiding the reader through a complete analysis of the freely available openfMRI ds000117 “familiar vs. unfamiliar vs. scrambled faces” dataset. The goal is to educate both novice and experienced data analysts with the “tricks of the trade” necessary to successfully perform this type of analysis on their own data.

## Introduction

In this paper, we demonstrate the application of dynamic imaging of coherent sources (DICS), a spatial filtering technique for magneto/electro-encephalography (MEG/EEG) data originally proposed by Gross et al. (2001). Spatial filters, or beamformers, are constructed to pass the activity originating at a specific location, while suppressing activity from other locations using a weighted sum of the sensor signals.^1^ DICS is a linearly constrained minimum variance beamformer in the frequency do-main, which can be used to calculate oscillatory power at any given location in the brain and coherence between any two given locations.^2^ This enables us to create cortical “power maps” and to perform functional connectivity analysis.

Interacting large-scale functional networks in the brain are thought to support cognition and behavior. Dynamic changes in connectivity are of increasing interest, as recent results have shown that functional connectivity between brain regions changes in a time-resolved and task-dependent manner^3^ and hence provides information that is complementary to the analysis of evoked responses.^4^ Magnetoencephalography (MEG) recordings provide a direct measure of neural activity with excellent time resolution. MEG enables non-invasive estimation of connectivity between brain regions with a cortex-wide spatial coverage that cannot be attained with, for example, intracranial recordings.

There are different ways to define and quantify functional connectivity.^5^ In general, two regions are assumed to interact when certain aspects of the recorded brain activity over these regions are consistent. In this paper, we focus on coherence, which quantifies the cortico-cortical synchrony of oscillatory activity, as a connectivity measure.^6^ Oscillatory activity in neuronal populations is a principal feature of brain activation and synchronization, or coherence, of such oscillating activity across brain regions is thought to promote efficient communication within large-scale neural networks.^7^ Coherence is thus a neurophysiologically well motivated measure of functional connectivity. Previous studies have suggested that oscillatory activity/interaction within specific frequency bands may have different functional roles.^8^ Using coherence as a measure of connectivity enables a direct mapping of connectivity at different frequencies, without the need to estimate time series at the level of cortical sources.^9^

Recently, we developed a pipeline for estimating all-to-all functional connectivity^10^ for MEG network analysis, which utilizes the DICS spatial filter combined with a wavelet approach to achieve a high temporal resolution.^11^ With this approach, we have demonstrated that a transient reorganization of the large-scale functional networks that support language takes place before onset of speech.^12^

For the current paper, we have made a new implementation of our pipeline and integrated it with the MNE-python package.^13^ We will demonstrate it using the freely available MEG dataset collected by Wakeman and Henson (2015), for which we have chosen to compare changes in oscillatory activity and functional connectivity between processing faces and scrambled images, as described in section 1.1. We will go over all the steps of the analysis and provide examples of how to implement them using MNE-Python.

The preprocessing of the MEG data is briefly outlined in section 2. In the DICS beam-former, a cross-spectral density (CSD) matrix is used to represent the measured oscillatory activity and their dependencies. In section 3, we describe estimation of the CSD matrices, the mathematical formulation, and its implementation using the python code. For group-level comparisons it is important to obtain comparable source-points and connections across subjects. For this purpose, we have chosen to create a surface-based cortical grid in a template brain and transform the source locations to each individual subject. In section 4, we outline how this is implemented.

While the current pipeline was primarily developed for the purpose of all-to-all connectivity analysis, it can also be used for estimation of oscillatory activity, (i.e., “power mapping”), which we discuss in section 5. In section 6, we introduce a “canonical” computation of coherence between brain regions, which facilitates the stable estimation of all-to-all connectivity.^14^ In this approach, the source orientation configuration for each cortico-cortical connection is determined by identifying the orientation combination that maximizes coherence between the two sources.

Neurophysiological recordings are inherently sensitive to spatial blurring of the signal due to field spread, thus complicating the estimation of functional connectivity between brain regions.^15^ Hence, we focus on connections that span long distances (>4 cm). To further suppress effects related to field spread, the current approach is based on identifying statistically significant differences in functional connectivity between power-matched experimental conditions, rather than absolute coherence values. This analysis step is described in section 7.

Importantly, this approach identifies changes in connectivity between brain regions that can be linked to the specific task manipulation, rather than the entire underlying network. For visualization of the identified networks, we use a combination of a cortical-level degree map which shows the total number of connections for each source point, and a circular connectogram that summarizes the number of connections between brain regions at a cortical parcellation level. This is presented in section 8.

Finally, we discuss benefits and limitations of the present approach in section 9, and present several methodological considerations related to functional connectivity analysis with MEG.

### Example dataset

Throughout this paper, we will demonstrate the application of our pipeline to an example dataset. For this purpose, we use the data collected by Wakeman and Henson (2015). This subsection will provide a brief description of the characteristics of the data that are most salient to the present paper. For further details on the dataset, see Wakeman and Henson (2015).

The dataset consists of simultaneous MEG and EEG recordings, collected from 19 participants who were viewing images of either faces or scrambled versions of the face stimuli. The original study excluded data from 3 participants due to the presence of artifacts in the data^16^; we also excluded those data from our analysis. The data was recorded by an Elekta Neuromag Vectorview 306 system that has 204 planar gradiometers. Only the gradiometer MEG data was used in our example.

The stimuli consisted of 300 greyscale photographs, half from famous people (known to the participants) and half from people unknown to participants, and 150 images of scrambled versions of either famous or unknown faces. Each stimulus was presented twice, for a total of 2 × (300 + 150) = 900 trials, with the second repetition occurring either immediately after the first, or with an interval of 5–15 intervening stimuli. In our example analysis, we focus on the distinction between faces versus scrambled images, regardless of whether the faces were known or unknown to the participant.

Each trial began with the presentation of a fixation cross for a random duration of 400 ms to 600 ms, followed by presentation of the stimulus for a random duration of 800 ms to 1000 ms, after which a white circle was presented for 1700 ms. The task for the participants was to press one of two buttons depending on whether they judged the image to be “more” or “less” symmetric than average.

### Data and code availability

The multi-subject, multi-modal human neuroimaging dataset^17^ that we use in this study can be found at: https://openfmri.org/dataset/ds000117.

The code repository related to this project is at: https://github.com/wmvanvliet/conpy. This currently includes the ConPy project code (in the `conpy/` folder), the analysis scripts to process the Wakeman and Henson (2015) dataset (in the `scripts/` folder), the scripts to produce the figures in this paper (also in the `scripts/` folder), the code examples included in this paper (in the `paper/code_snippets/` folder), and the 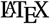 code to produce the final pdf (in the `paper/` folder). Further instructions on how to run the pipeline are provided in the `README.md` file.

## Preprocessing

DICS based power analysis and functional connectivity can be investigated for multiple kinds of experimental designs, ranging from ones consisting of isolated events^18^ to ones with continuous naturalistic stimulation.^19^ The present analysis pipeline focuses on data representing neural processes related to external stimuli or events whose timing can be determined exactly. For this type of analysis, an important preprocessing step is to cut up the continuous MEG recording into fragments of data surrounding the onset of such events. These fragments are referred to as “epochs”.

The process of going from raw data to epochs is not specific to DICS analysis, but are the first steps shared by many analysis pipelines. A sister paper, Jas et al. (2017), discusses the many parameters and trade-offs involved in these important preprocessing steps and provides code examples, using the same dataset as in this paper. Therefore, to keep the topic of the current paper focused on estimating cortical power and connectivity analysis, and to avoid duplication of effort, we refer the reader to Jas et al. (2017) for a detailed description of the preprocessing steps, which we will only summarize below.

Construction of the source space and forward model (see section 4) depends on a 3D model of the subject’s head that is created from a structural magnetic resonance imaging (MRI) scan, which is done in our analysis pipeline using the FreeSurfer^20^ package.

The MEG data is processed with the `maxfilter` program, developed by Electa and also implemented in MNE-Python, to eliminate noise sources that originate outside the MEG helmet. Furthermore, the program uses the head coils that are attached to the participant’s head to track the head position during the recording, and projects the data such that the influence of head movements is minimized.

To remove the signals produced by the head coils, the MEG signal must be low-pass filtered to at least below 150 Hz. Additionally, the signal should be high-pass filtered above at least 1 Hz when performing independent component analysis (ICA).

To reduce the contamination of the MEG signal by artifacts caused by eye blinks and heart beats, ICA components are estimated on the continuous data. However, no actual data decomposition is performed yet. Next, an automated detection algorithm is applied to detect the onset of blinks and heart beats, and segments of data surrounding each onset are created and averaged, yielding an “average blink” segment and average “heart beat” segment. The average blink and average heart beat segments are then decomposed along the ICA components and the correlation between the electro-oculography (EOG) and electro-cardiography (ECG) sensors and each signal component is computed. The ICA components for which the corresponding signal components correlate strongly with the EOG or ECG signal are flagged as “bad” and will be removed in the next step.

The continuous data is cut up into segments in a short time window relative to the onset of the presentation of each stimulus. These segments are referred to as “epochs”. The data of each epoch is decomposed along the ICA components that were computed in the previous step, the components flagged as “bad” are dropped, and the signal is recomposed. Finally, epochs where the signal amplitude of one or more channels exceeds a predefined threshold, signifying the presence of an artifact (for example such as those caused by movements and biting) that contaminates the data segment beyond repair, are removed.

### Application to the example dataset

For our analysis of the Wakeman and Henson (2015) dataset, we mostly follow the preprocessing pipeline of Jas et al. (2017), which implementation can be found at https://github.com/mne-tools/mne-biomag-group-demo. However, there are some key differences between our pipeline and the one used by Jas et al. (2017):

1. Since our pipeline operates on MEG data, we restrict our preprocessing pipeline to only use the gradiometer and magnetometer channels, whereas Jas et al. (2017) also include EEG data.
2. Since we are analyzing oscillatory activity rather than evoked potentials, we high-pass filter the data above 1 Hz, whereas Jas et al. (2017) perform no high-pass filtering other than the one performed by the recording hardware.
3. We do not make use of the `autoreject` package for dynamically determining thresholds when rejecting epochs which have a too large signal amplitude, but rather use a fixed, more lenient, threshold. This is because since our analysis of oscillatory activity is less sensitive to isolated signal spikes than the analysis of evoked potentials performed in Jas et al. (2017).

The preprocessing pipeline is implemented in the following scripts:

| Script | Function |
| --- | --- |
| 00_fetch_data.py | downloads the openfmri ds117 dataset and extracts it |
| 01_anatomy.py | runs the FreeSurfer recon-all program on the anatomical MRI data and uses the FLASH images to create surface meshes |
| 02_filter.py | performs band-pass filtering between 1 Hz to 40 Hz on the MEG data |
| 03_ica.py | uses ICA to decompose the MEG signal into independent components. Finds at most two ICA components that correlate with heart beats ($\rho > 0.05$), at most two components that correlate with eye blinks ($\rho > 0.1$), and flags these components for removal. |
| 04_epochs.py | cuts the continuous MEG data into epochs from $-0.2$ s to $2.9$ s relative to the onset of the stimuli. Removes ICA components that were flagged in the previous step. Removes epochs where the signal amplitude of one or more gradiometer channels exceed $3 \times 10^{-10}$ T/cm or one or more magnetometer channels exceed $4 \times 10^{-12}$ T. |

From this point on, the analysis pipeline becomes specific to DICS analysis and will be described in more detail.

## Estimating cross-spectral density (CSD) matrices

Estimating the cortical origins of oscillatory activity (we refer to this as “power mapping”) and estimating connectivity between cortical sources (we refer to this as “connectivity analysis”) both start with the computation of one or more cross-spectral density (CSD) matrices. The CSD is the covariance between the two signals, in our case the activity recorded at two sensors, in the frequency domain. A CSD matrix defines the CSD between all sensor-pairs and is similar in nature to a covariance matrix.

Commonly, both the analysis of oscillatory power and connectivity are conducted in multiple frequency bands, time windows, and/or experimental conditions. For each of these, a separate CSD matrix needs to be computed.

Because we wish to compute CSD matrices for specific frequency bands and time windows, we choose to transform the signal to the time-frequency domain using a wavelet transform. We follow the method outlined in Tallon-Baudry, Bertrand, Delpuech, and Permier (1997), which offers a better tradeoff between time and frequency resolution than a standard Fourier transform.

### Mathematical formulation

For each frequency *f* (in Hertz) we want to include in the analysis, we construct the corresponding Morlet wavelet **m**(*f*), which has a Gaussian shape both in the temporal and frequency domain. The standard deviation of this Gaussian shape in the time domain, *σ*_t_, is an important parameter that determines the tradeoff between temporal and frequency resolution of the resulting time-frequency decomposition. A common tactic is to use a large *σ*_t_ at low frequencies, increasing the frequency resolution at the cost of temporal resolution, and use increasingly smaller values at higher frequencies, trading frequency resolution for temporal resolution. A convenient way to achieve this is to define *n*_o_ as the number of oscillations the Morlet wavelet completes. Then,

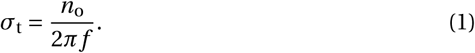

A Morlet wavelet of the desired length can then be constructed as follows:

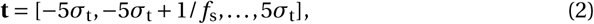

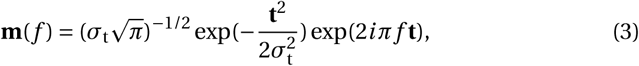
where **t** are the time points at which the Morlet function is evaluated and *f*_s_ is the sampling frequency of the MEG signal. The transformation to the time-frequency domain is performed by convolution of the Morlet wavelet for each frequency with the MEG signal:

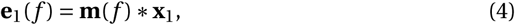

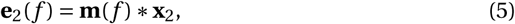
where (*) denotes linear convolution. The resulting vectors **e**_1_(*f*) and **e**_2_(*f*) contain the complex time courses of the signals **x**_1_ and **x**_2_ filtered at frequency *f*. Finally, we compute the CSD between the signals by taking the dot product of **e**_1_(*f*) and the complex conjugate of **e**_2_(*f*) for all the frequencies and time points we wish to include in the analysis and by averaging the result:

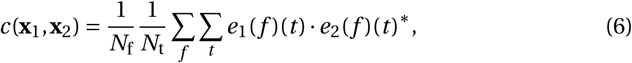
where *N*_f_ is the number of frequencies, *N*_t_ the number of time points, (·) denotes the dot product between two vectors, the superscript (*) the complex conjugate operation and *e*(*f*)(*t*) the signal at frequency *f* and time *t*. Since the frequency domain is described using complex numbers, *c* will be a complex number as well when the computations are done for distinct signals **x**_1_ and **x**_2_.

To compute the full CSD matrix, equation 6 is repeated for each pair of channels. Each element *C*(*i*, *j*) of the resulting CSD matrix 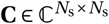 holds the CSD between sensors *i* and *j*. The matrix is Hermitian, so *C* (*i*, *j*) and *C* (*j*, *i*) are complex conjugates of each other, and the diagonal elements hold the mean power-spectral density (PSD) for each sensor. The CSD matrices are computed for each epoch separately and then averaged to produce a single CSD matrix per experimental condition.

### Code example

The following code example will compute the CSD matrix over the time range from 0 s to 0.4 s relative to the stimulus onset, for two frequency ranges:

~~~
**import** numpy as np, mne                              # Import required Python modules
epochs = mne.read_epochs(‘sub002-epo.fif’)       #Read epochs from FIFF file
epochs = epochs[‘face’]                              #Select the experimental condition
frequencies = np.linspace(7, 17, num=11)         #Specify frequencies to use
csd = mne.time_frequency.csd_morlet(epochs, frequencies, tmin=0, tmax=0.4, n_cycles=7, decim=20)
csd_alpha = csd.mean(7, 13)                         # CSD for alpha band: 7–13 Hz
csd_beta = csd.mean(13, 17)                         # CSD for beta band: 13–17 Hz
~~~

As discrete wavelet transforms are used in the CSD computation, the frequencies are specified as a list, rather than a range. These frequencies should evenly span the desired frequency range. Their suitable spacing depends on the frequency resolution of the wavelets.

The `n_cycles` parameters of the `csd_epochs` function controls *n*_o_, thus controlling the tradeoff between frequency and time resolution of the wavelet transform. It can either be set to a fixed value (as in the example), which means the wavelets get shorter as the frequency increases (increasing the temporal resolution and decreasing the frequency resolution). Alternatively, one may specify a list of values, one for each frequency, to have precise control over the time/frequency resolution tradeoff.

The `decim` parameter of the `csd_epochs` function controls the spacing of the time points *t* that are used in equation 6, enabling more efficient computation of the CSD matrix. The time resolution of the signals following the wavelet transform (equation 4) is generally much lower than the sampling rate of the original signals. In these cases we can safely pick every n^th^ time point without losing information.

The wavelet convolution method assumes that the data across time has an approximate mean of zero. In our pipeline, we choose to remove the signal offset for each epoch. Another good option is to first apply a highpass filter to the data, in which case further detrending is not necessary.

### Application to the example dataset

For our analysis of the Wakeman and Henson (2015) dataset, we computed CSD matrices for the following frequency bands (following Liljeström, Kujala, et al. (2015)):

| Frequency range | Band name |
| --- | --- |
| 3 Hz to 7 Hz | theta |
| 7 Hz to 13 Hz | alpha |
| 13 Hz to 17 Hz | low beta |
| 17 Hz to 25 Hz | high beta 1 |
| 25 Hz to 31 Hz | high beta 2 |
| 31 Hz to 40 Hz | low gamma |

We choose to use a fixed *n*_o_ = 7, and the width of our chosen frequency bands reflect the resulting time/frequency resolution tradeoff. There is no golden standard for which frequency bands to use and you may have to adapt the frequency ranges to fit your dataset and research question. For example, frequencies higher than 40 Hz may be of interest as well.

The CSD matrices were computed for both the time window from 0 s to 0.4 s, and during the “baseline” period from −0.2 s to 0 s, relative to the onset of the stimulus. We will later compare the cortical sources of oscillatory activity before and after the presentation of a stimulus. This analysis step is implemented in script `05_csd.py` and an example of the resulting CSD matrices is presented in figure 1.

**Figure 1:**
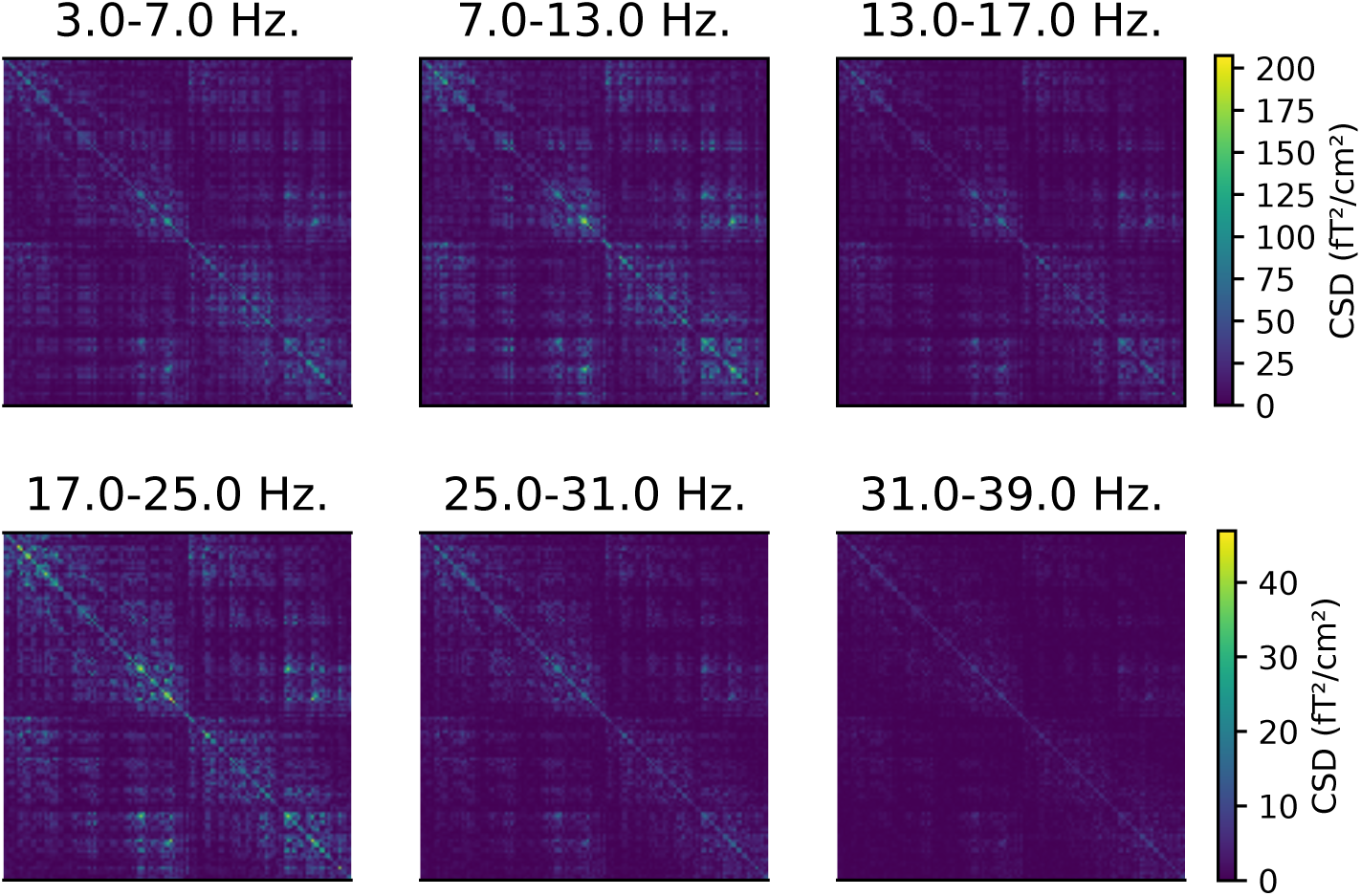
CSD matrices computed for different frequency bands. The CSD matrices were computed across all the epochs where a face stimulus was presented to subject 2, in the time window from 0 s to 0.4 s relative to the presentation of the stimulus. Each row and column corresponds to one of the 204 gradiometers. Note that each row has a separate color scale.

## Source space and forward model

The DICS beamformer will, given a CSD matrix and forward modeling of neural currents, estimate the power of the oscillatory activity originating from one specific point on the cortex. DICS uses a spatial filter to determine the activity at the given point on the cortex while suppressing contributions from all other sources. By creating a grid of regularly spaced points along the cortex and computing spatial filters for each point, a complete picture of brain-wide activity emerges. This grid is referred to as the “source space” (figure 2, left). The DICS power estimates are also used during connectivity analysis, where the source space is used for defining the start and end point of possible connections.

**Figure 2:**
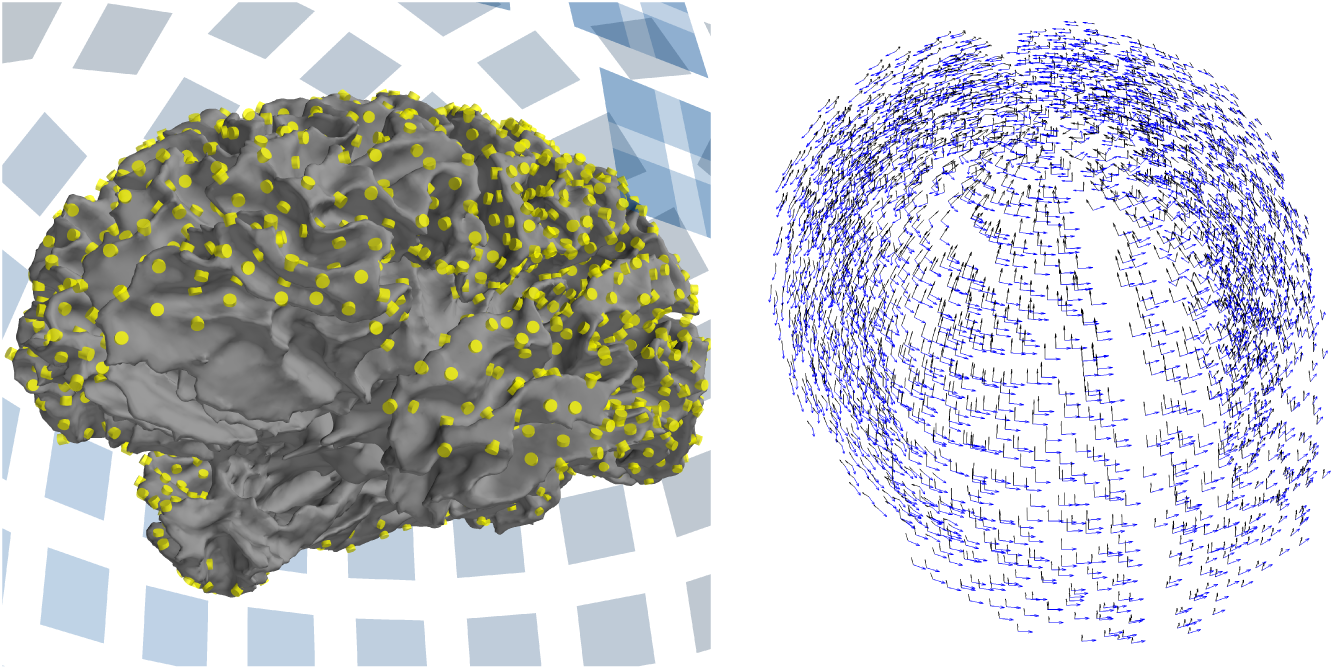
The source space and forward model used in connectivity analysis. **Left:** The white matter surface, as reconstructed by FreeSurfer. The source space is defined as a grid of points along this surface, shown in yellow. All points further than 7 cm from the closest MEG sensor (shown as blue squares in the background) have been discarded. **Right:** The forward model defines two dipoles at each source point. The orientation of the dipoles is tangential to a sphere with its origin at the center of the brain.

To create the source space, we first need a 3D-model of the subject’s brain. Here, we obtain it by performing a structural MRI scan on the subject and processing the data with FreeSurfer.^21^ The details are explained in Jas et al. (2017) and the implementation can be found in the script `01-run_anatomy.ipy` accompanying that paper. The FreeSurfer analysis results in several 3D meshes, corresponding to different brain tissues, of which the white matter surface serves as the basis for our source space.

For group-level analysis, it is important that connections between the points within source spaces can be compared across subjects. This is feasible if the same connections exist for each subject, which, in turn, means that the same source points must be defined for each subject. To facilitate this, we first define the source space on the “fsaverage” brain: a template brain model, provided by FreeSurfer, constructed by averaging the MRI scans of 40 subjects.^22^ The resulting source space is then morphed to each individual subject, transforming the source points to corresponding locations on the cortex.^23^ Note that the morphed source space will generally be only approximately evenly spaced. For creating power maps, we advice to create evenly spaced source spaces for each individual subject and morph the estimated power map to the average brain, as explained in Jas et al. (2017).

In our analysis pipeline, we compute all-to-all connectivity between the source points. To keep the number of connections manageable, only a limited number of source points can be used. Partly, this is facilitated by placing the sources at slightly larger spatial intervals than is common in studies focusing on cortical activity. In addition, we place sources only in areas that can be reliably measured using MEG, rather than in “deep sources” that do not generate signals that would be readily detectable with MEG sensors.

We start out with a regularly spaced grid of 5124 points covering the entire surface of the cortex, yielding an average distance of 2.6 mm between neighbouring points. To limit the number of source points, all points that are further than 7 cm from the nearest MEG sensor are discarded. For this dataset, a cutoff distance of 7 cm provides a good tradeoff between the number of source points and coverage across the cortex, but this value may need to be adjusted for other datasets. Close visual inspection of the result is required, see figure 2 (left). To ensure that the same source points are defined for each subject, the distance from source points to the closest sensor is determined in one subject, and the resulting set of points is then used for all subjects. Since the distance from the source points to the sensors is dependent on the position of the subject’s head in the MEG helmet, it is important to ensure that the initial distance computations are done for a subject whose head was in an approximately average position across subjects with respect to the helmet.

The resulting restricted source spaces are only used during connectivity analysis. For computing power maps, the number of source points is less of an issue and therefore we always use the full source space.

Given the source space, we construct a forward model that models how the magnetic field, produced by a current at each source point, travels through the various tissues of the brain and head, resulting in activity recorded at the MEG sensors. For this computation, we employ a boundary element method (BEM) model^24^ that uses the FreeSurfer meshes of the brain tissues, assuming homogeneous conductivity within each mesh. For MEG datasets, we only include the inner skull meshes, resulting in a single-layer BEM model.

The neural currents at the source points are modeled as equivalent current dipoles (ECDs) that represent the dominant component of the local current as a vector that has both a magnitude and a direction. The forward model represents the ECD at each source point using three separate dipoles, arranged in three orthogonal orientations, representing the magnitude of the current in the x-, y-, and z-directions. We will refer to these orthogonal dipoles, which are merely mathematical constructs, simply as “dipoles”, while we will refer to the source dipole that is formed by combining the three orthogonal dipoles, as “the ECD”.

During the connectivity computation, we reduce the number of dipoles for computational efficiency reasons (section 6.1) and use only two orthogonal dipoles instead of three; specifically, we use two orthogonal dipoles that are tangential to a spherical approximation of the head shape (figure 2, right) and that generate stronger magnetic fields than radial sources.^25^ For computing power maps, we prefer to use three orthogonal dipoles at each source point.

### Code example

The following code example will construct a forward model for a single subject, suitable for connectivity analysis, following all the steps outlined above:

~~~
**import** conpy, mne # Import required Python modules
~~~

~~~
# Define source space on average brain, morph to subject
src_avg = mne.setup_source_space(‘fsaverage’, spacing=‘ico4’)
src_sub = mne.morph_source_spaces(src_avg, subject=‘sub002’)
~~~

~~~
# Discard deep sources
info = mne.io.read_info(‘sub002-epo.fif’) # Read information about the sensors
verts = conpy.select_vertices_in_sensor_range(src_sub, dist=0.07, info=info)
src_sub = conpy.restrict_src_to_vertices(src_sub, verts)
~~~

~~~
# Create a one-layer BEM model
bem_model = mne.make_bem_model(‘sub002’, ico=4, conductivity=(0.3,))
bem = mne.make_bem_solution(bem_model)
~~~

~~~
# Make the forward model
trans = ‘sub002-trans.fif’ # File containing the MRI<->Head transformation
fwd = mne.make_forward_solution(info, trans, src_sub, bem, meg=True, eeg=False)
~~~

~~~
# Only retain orientations tangential to a sphere approximation of the head fwd = conpy.forward_to_tangential(fwd)
~~~

For group-level analyses, it is important to note that MNE-Python stores the source points as vertex indices of the original FreeSurfer mesh and that these indices are always stored in sequential order. Thus, when we morph the source space defined on the “fsaverage” brain to an individual subject, the ordering of the source points is not preserved. For example, the first source point of subject 1 can correspond to the fourth source point of subject 2. To account for this, we always store vertex indices in the order defined in the “fsaverage” source space. To re-order the individual-level source-points correctly, we first determine the changes in the ordering of the vertices using the `conpy.utils.get_morph_src_mapping` function and modify the vertex indices accordingly. This process is implemented in script `07_forward.py`.

### Application to the example dataset

In the example dataset, the source space was first defined on the “fsaverage” brain and then morphed to each subject. For each subject, three orthogonal dipoles were placed at each source point and the white matter and skull FreeSurfer meshes were used to compute the forward model. The construction of the source spaces for the “fsaverage” brain is implemented in script `06_fsaverage_src.py` and the morphing of the source space to the brains of the individual subjects and subsequent computation of the forward models are implemented in script `07_forward.py`.

While power mapping was done using all the source points, connectivity analysis used a restricted source space where all source points further than 7 cm from the closest MEG gradiometer were discarded. This distance measurement was performed on the first subject and then used for all other subjects. This process is implemented in script `08_select_vertices.py`. In connectivity analysis, the forward models that define three dipoles at each source point were transformed into “tangential” models that define two dipoles at each point. This step is implemented in script `10_connectivity.py`.

### Power mapping

The DICS beamformer can be used to estimate the cortical sources of oscillatory activity within a given frequency band. As explained in section 4, a grid of source points is defined along the cortex. At each source point, three current dipoles are defined that are arranged to have orthogonal orientations. A whole-brain estimate of the oscillatory power is produced by computing, for each dipole, a spatial filter that passes activity that can be attributed to the dipole, while reducing activity originating from other sources.^26^

Other than the various parameters involved in computing the CSD matrix and the forward model, the “regularization” parameter is an important parameter governing the creation of the spatial filters. In practice, the regularization parameter represents a tradeoff between the amount of detail in the power maps and their sensitivity to noise. If the amount of regularization is too small, it may result in the estimates being driven by noise factors, yielding sub-optimal results. If too much regularization is used, relevant details may be obscured and the power map will be dominated by the strongest sources. Typical values are in the range 0.01–0.1, scaled by the mean singular values of the CSD matrix.

The resulting cortical power maps define, at each source point, the power in all orientations. Typically, for each source point, only the power corresponding to the orientation that maximizes the power is reported.

### Mathematical formulation

The regularization parameter *α* arises from the need to compute the inverse of the CSD matrix. Since this matrix is often rank deficient, its inverse cannot be directly computed, but a pseudo-inverse needs to be approximated. This estimation is more stable when a small value is added to the diagonal (diagonal loading):

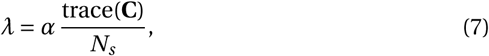

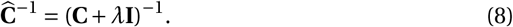

We use the Moore–Penrose pseudoinverse to compute (**C** + *λ***I**)^−1^.

Initially, the power maps will be biased towards superficial sources, since they have a larger effect on the MEG sensors. To counter this, the leadfields can be normalized before computing the spatial filters:

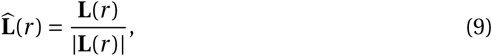
where **L**(*r*) is a row vector containing the leadfield connecting dipole *r* to each sensor, |·| denotes the norm of the vector and 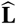 is the normalized leadfield.

The DICS beamformer is a linearly constrained minimum variance (LCMV) beam-former, computed using and applied to a (time-)frequency transformation of the original signals. We deviate slightly from Gross et al. (2001) by computing the filter for each dipole separately:

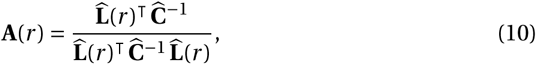
where **A**(*r*) is a vector of weights that constitutes a linear spatial filter that attempts to isolate the signal power for the dipole from the rest of the signal. In our approach, we treat dipoles with different orientations as separate sources, even if their locations are the same, and consequently compute the beamformer filter for each dipole individually. In this case, 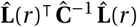 reduces to a scalar value, which avoids having to compute the inverse of another rank deficient matrix. We obtain an estimate of the power at a source point by multiplying the filters for all dipoles defined at the location with the CSD matrix:

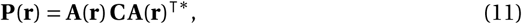
where **A**(**r**) is a matrix whose rows contain the filters for all dipoles **r** defined at the source point and **P**(**r**) is the resulting power estimate. The power estimate contains, along the diagonal, the square of the power at each dipole, and the off-diagonal elements contain the cross-power estimates between dipoles.

Common methods of summarizing **P**(**r**) are:

1. choosing the direction that maximizes the power, i.e. the first singular value of **P**(**r**)
2. the sum of the squared power for each dipole, i.e. trace(**P**(**r**))
3. the squared power in the direction that is orthogonal to the surface of the cortex.

### Code example

In the following example, we compute the cortical power maps for oscillatory activity in the range from 7 Hz to 13 Hz for the epochs corresponding to trials where a face stimulus was presented:

~~~
**import** mne # Import required Python modules
info = mne.io.read_info(‘sub002-epo.h5’) # Read info structure
fwd = mne.read_forward_solution(‘sub002-fwd.fif’) # Read forward model
csd = mne.time_frequency.read_csd(‘sub002-csd-face.h5’) # Read CSD
csd = csd.mean(fmin=7, fmax=13) # Obtain CSD for frequency band 7–13 Hz.
# Compute DICS beamformer filters
filters = mne.beamformer.make_dics(info, fwd, csd, reg=0.05,
                                                           pick_ori=‘max-power’)
# Compute the power map
stc = mne.beamformer.apply_dics_csd(csd, filters)
~~~

The regularization parameter `reg` is set here to 0.05, which is generally a good tradeoff between the level of spatial detail and sensitivity to noise. It is good practice to experiment with different values to see how the power maps behave: if the power estimates change substantially for small increments of the `reg` parameter, it may be set too low. The `pick_ori` parameter selects the method with which to summarize the power at each source point. In this case, for each source point, the power is computed along the direction which maximizes the power.

When comparing the power maps from different subjects, the `stc` objects can be morphed to the “fsaverage” brain with the `stc.morph(to_subject)` method. The morphed `stc` objects can then be straightforwardly averaged and analyzed using the statistical functions in the same manner as for other types of source estimates.^27^

### Application to the example dataset

The scripts `09_power.py` and `11_grand_average_power.py` implement the full analysis on the example dataset. Script `figure_power.py` produces figure 3.

**Figure 3:**
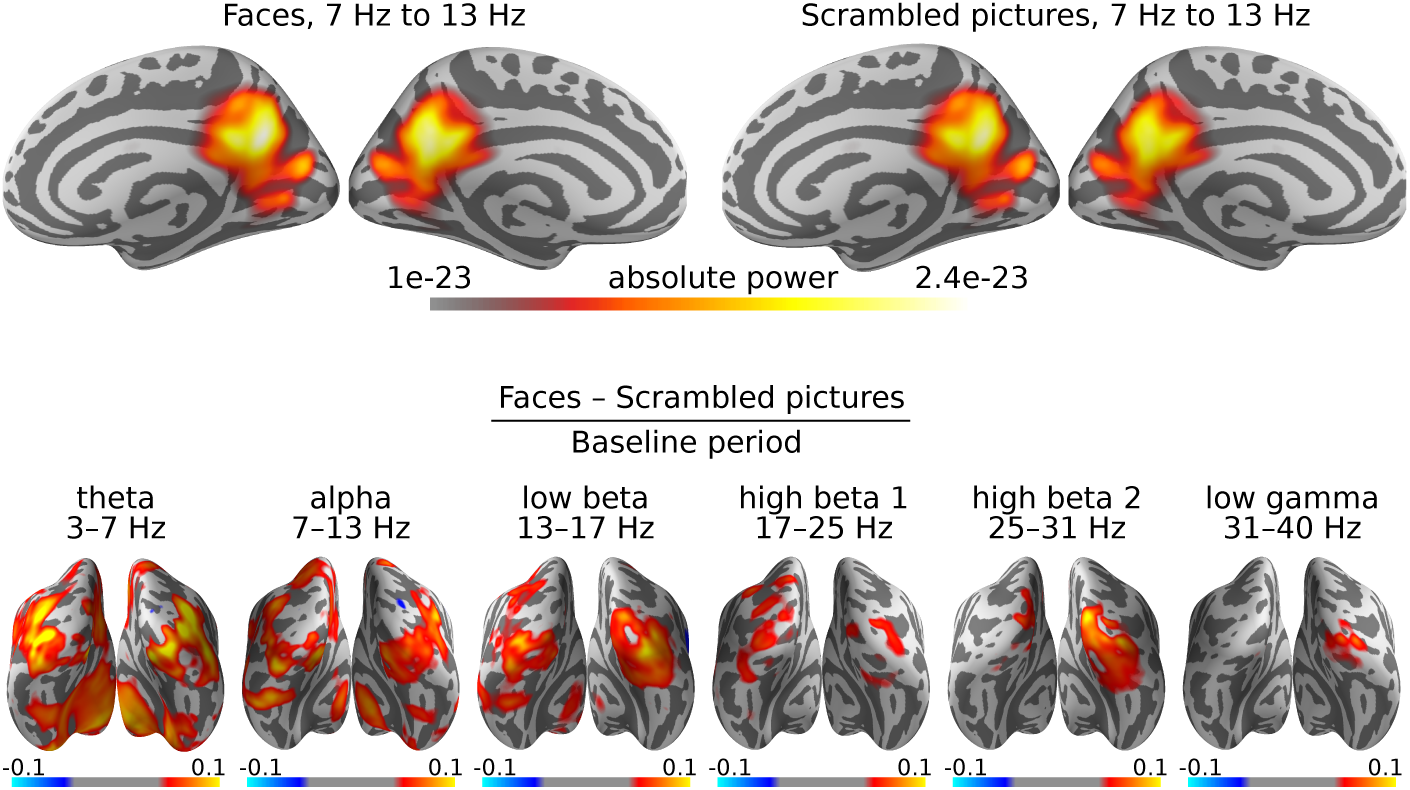
DICS grand average power maps. Cortical activity is visualized on an “inflated” version of the cortex, so as not to hide activity within the sulci. **Top:** Estimation of cortical origins of oscillatory activity in the alpha band. In this case, the inflated view makes it seem there are three sources of alpha power, but in reality, these sources are adjacent on the original white matter surface. **Bottom:** Contrasts between faces and scrambled images for all frequency bands. Warm colors indicate sources with more activity for faces than scrambled images and cold colors indicate sources with less.

It is common for the power maps to be dominated by alpha and/or beta activity, as is the case for our example dataset as well (figure 3, top row). The alpha rhythm is typically generated in the parieto-occipital cortex. The beamformer localizes the alpha activity over the entire 0 s to 0.4 s time window as a single, somewhat deep source.^28^

More interesting effects are revealed by contrasting two experimental conditions. In the case of the example dataset, these are the presentation of faces versus scrambled images. Furthermore, we are interested in the changes in oscillatory power caused by the presentation of the stimuli, relative to the baseline period. Accordingly, our final power maps are computed as “(faces−scrambled pictures)/baseline” (figure 3, bottom row).

The experimental paradigm used in the example dataset was designed to produce strong evoked potentials (EPs). Although DICS aims to capture oscillatory activity, the power maps are likely dominated by the EPs, especially in the lower frequency bands. In our case, all frequency bands highlight the primary visual cortex, where there is a strong EP following shortly after the presentation of a visual stimulus. The lower-and upper-beta band also show activity in the fusiform face area in the right hemisphere that is more active during face stimuli than scrambled images. The theta and alpha bands show some frontal power differences that are likely due to residual eye movement artifacts that were not completely removed in the ICA step. The low gamma frequency band only shows some very slight increases in activity, which is why we chose to perform the connectivity analysis for this frequency band, since large differences in power between conditions will severely bias an all-to-all connectivity estimate.

For better interpretation of these results, one can proceed with statistical analysis of the power maps in a similar fashion as done with source estimates of evoked data, as detailed in Jas et al. (2017).

## Connectivity analysis

In addition to analysis of oscillatory power, DICS is commonly used to investigate connectivity between cortical areas. The DICS beamformer is well suited for estimating cortical connectivity, as coherence between brain regions can be determined based on the sensor-level CSD matrices, without the need to first estimate the time courses for the regions of interest, as required for most other connectivity metrics. The coherence metric quantifies the level of synchronicity between the oscillatory activity of different areas, on a scale from 0 (no synchronization) to 1 (perfectly synchronized). Coherence is thought to be indicative of inter-areal communication.^29^

Ideally, one would compute coherence between all source points in the source space. However, in practice, this is currently computationally intractable, so several thresholds will be applied to prune the number of connections. In section 4, the first threshold was applied, namely that deep sources were eliminated from the source space. This has the effect of only considering source locations where the MEG signals are the most reliable. The second threshold we apply is a distance criterion. Due to the inherent field spread of the MEG signal,^30^ source points that are close together will always exhibit strong coherence. While this effect is alleviated by considering a contrast between two conditions, long-range connections^31^ can be estimated more reliably than short-range ones. For this reason, all connections between source points which are closer than a distance threshold (e.g., ≤4 cm) are removed from further analysis. The distance threshold is a parameter that needs to be chosen with care and in consideration with the research question of the study. When interpreting the result, one should always remember that there may be additional short-range connections present, but hidden from view due to the distance threshold.

In order to perform group-level analysis, coherence must be computed for the same connections in each subject. Therefore, the distance threshold based pruning is first applied to the connectivity pairs in a single subject, and the selection is subsequently carried over to the other subjects. In section 7, connections are further pruned based on a contrast between the experimental conditions.

### Canonical computation of coherence

The connectivity computation is complicated somewhat by the fact that, generally, the forward model defines currents with both a magnitude and an orientation, represented through the use of multiple dipoles at each source point. For example, our connectivity pipeline employs a “tangential” forward model that defines two orthogonal dipoles tangential to a sphere (see section 4). As mentioned in the section on power mapping (section 5), there are several ways to summarize the information at each source point. One way would be to only use the orientation that maximizes source power.^32^ However, simulations that were performed as part of the study by Saarinen et al. (2015) have shown that this strategy tends to produce spurious increases in coherence between unsynchronized sources. A better strategy may be to use, for each connection, orientations that maximize the coherence between the two source points. This involves going through all the possible orientation combinations for the two source points, and choosing the orientation pair that maximizes the coherence. We refer to this strategy as “canonical computation of coherence” and it is the default strategy implemented in the ConPy package.

### Mathematical formulation

Given a CSD matrix **C**, it is straightforward to compute coherence between sensors (and later between cortical regions). The coherence *m* between sensors *i* and *j* is:

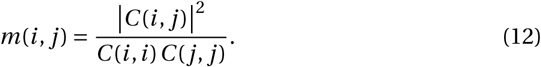

To compute coherence between source points, the CSD matrix is first run through the DICS beamformer to obtain power estimates at each source point. In our canonical coherence pipeline, we deviate from Gross et al. (2001) and replace the CSD matrix **C** in equation 11 by the regularized version 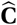. This results in an approximation of the power that is much faster to compute, as the equation simplifies to:

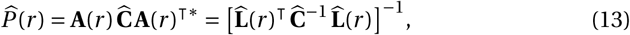
where 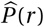 is an approximation of the power estimate for dipole *r*. Similarly, the cross-power between two dipoles (*r*_1_, *r*_2_) is approximated by 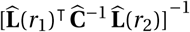.

In the canonical computation of coherence, coherence is estimated by optimizing the orientation of the ECDs at both source points for each connection. Here, we employ a tangential forward model, which defines two orthogonal dipoles at each source point to encode information about the leadfield in different orientations. Using the tangential source orientation plane, we denote the leadfield for an ECD with orientation *θ* as:

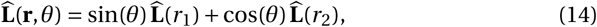
where *r*_1_ and *r*_2_ are the two dipoles defined at the source point and **r** = [*r*_1_, *r*_2_].

Canonical coherence between two source points *M*(**r**_1_, **r**_2_) is computed as follows:

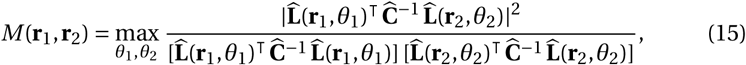
where **r**_1_ are the two dipoles defined at the first source point and **r**_2_ are the dipoles defined at the second source point, and *θ*_1_ is the orientation of the ECD at the first source point and *θ*_2_ the orientation of the ECD at the second source point.

The computation max_*θ*_1_,*θ*_2__ is conducted by performing a search over all possible ECD orientation combinations and using the maximum coherence value encountered during the search. In practice, ca. 50 different orientations are evaluated at both locations, spanning the tangential orientation plane at discrete intervals.

### Code example

In the following example, we compute connectivity between all combinations of source pairs that are at least 4 cm apart. For each connection, we compute the coherence between ECDs that are oriented in such a manner that the coherence between them is maximized (canonical computation of coherence). To reduce the search space for the optimal orientation, we convert the forward model from one with three dipoles at each source point, to a tangential model with two (figure 2, right), which limits the orientations to the tangential plane:

~~~
**import** conpy, mne # Import required Python modules
# Read and convert a forward model to one that defines two orthogonal dipoles
# at each source, that are tangential to a sphere.
fwd = mne.read_forward_solution(‘sub002-fwd.fif’) # Read forward model
fwd_tan = conpy.forward_to_tangential(fwd_r) # Convert to tangential model
~~~

~~~
# Pairs for which to compute connectivity. Use a distance threshold of 4 cm.
pairs = conpy.all_to_all_connectivity_pairs(fwd_tan, min_dist=0.04)
~~~

~~~
# Load CSD matrix
csd = conpy.read_csd(‘sub002-csd-face.h5’) # Read CSD for ‘face’ condition
csd = csd.mean(fmin=31, fmax=40) # Obtain CSD for frequency band 31–40 Hz.
~~~

~~~
# Compute source connectivity using DICS. Try 50 orientations for each source
# point to find the orientation that maximizes coherence.
con = conpy.dics_connectivity(pairs, fwd_tan, csd, reg=0.05, n_angles=50)
~~~

When performing group-level analysis, it is important that connectivity is evaluated between the same pairs of source points in the same order across subjects. However, as we saw in section 4.1, by default, the ordering of the source points differs between subjects. Therefore, before comparing coherence values across subjects, the connectivity estimates need to be transformed to define the source points in the same order, e.g., the order of the “fsaverage” brain, with the `con.to_original_src` method.

### Application to the example dataset

For the example dataset, connectivity was estimated for the low gamma frequency band (31 Hz to 40 Hz) in each subject. The connectivity pairs were computed for the first subject and then carried over to the other subjects. This computation is implemented in script `08_select_vertices.py`. The connectivity computations are implemented in script `10_connectivity.py`. The visualization of the connectivity results is performed after computing group-level statistics.

## Group-level statistics

Our analysis pipeline is designed for studying changes in cortico-cortical connectivity between different experimental conditions (as opposed to resting state analysis which studies the naturally occurring network while the subject is “at rest” in the scanner^33^). Thus, instead of attempting to map the entire network, we focus on the parts of the network where connectivity changes between experimental conditions. This means that the experimental design plays a vital role in our analysis pipeline, as experimental conditions must be designed so that contrasting them will reveal the sub-network of interest and are power-matched to minimize the effects of field spread.

All-to-all connectivity results can give an overwhelming amount of information that can be difficult to interpret. One way to manage the complexity is to compute connectivity between parcels, rather than source points. However, in this paper we will demonstrate an alternative approach that focuses on pruning connections until a manageable number remains. The procedure is an adaptation of the non-parametric cluster-permutation test by Maris and Oostenveld (2007), where the difference is in the way the data is clustered.

Starting from the initial all-to-all connectivity estimate, we prune connections that do not show a reliable difference between the experimental conditions. To this end, we perform a paired *t* -test for each connection, comparing the coherence values for all subjects between the conditions. All connections with an associated absolute *t*-value below a given threshold are pruned, while the surviving connections are grouped into “bundles”. A “bundle” means in this context a group of connections whose start and end points are in close proximity to each other. Bundles can be found by constructing a six-dimensional space, where each connection is assigned a position based on the Cartesian (xyz) coordinates of its starting and end points, and performing a hierarchical clustering in this space. This clustering procedure is performed separately on connections with positive vs. negative *t*-values, to assure that a bundle only contains connections that have an experimental effect in the same direction. Each bundle is assigned a “bundle-*t*-value” by summing the absolute *t*-values of the connections inside the bundle.

To determine which bundles show a significant effect, we repeat the above procedure many times with randomly permuted data to model the distribution of bundle-*t* values we may expect from random data. Random data was produced by flipping the condition labels for a random number of subjects, choosing a new random set of subjects for each permutation. Importantly, for each random permutation, only the maximum bundle-*t*-value is appended to the list of randomly observed *t*-values. This is an effective way to manage type-I errors.^34^ Any bundle with a bundle-*t*-value that is higher than at least 95 % of the randomly obtained bundle-*t*-values, is deemed significant (*p* ≤ 0.05).

This procedure has two important parameters: the initial *t*-value threshold (`cluster_threshold`) for pruning connections and the maximum distance between connections to be considered part of the same bundle (`max_spread`). Both parameters have an effect on the size of the bundles and hence the sensitivity of the test. Since the bundle-*t*-values are the sum of the *t*-values of the individual connections, large bundles will usually have a large bundle-*t*-value, making them more likely to survive the statistical threshold. However, the cluster-permutation test only tells whether a bundle as a whole is significant, not which connections inside a bundle drive this significance. This means that a bundle that was flagged as significant could contain many connections that show little difference between experimental conditions, as long as it also contains connections that do show a salient difference.

In practice, we advise choosing `cluster_threshold` such that a manageable number of connections remain (up to a few thousand) and `max_spread` such that a reasonable number of connections (tens to hundreds) are assigned to each bundle. When choosing these parameters, it may help to visualize the selected connections (see section 8) before performing the permutation test.

### Code example

The following example reads in the connectivity objects for all subjects and all conditions and prunes the connections using the statistical thresholds outlined above.

~~~
**import** conpy, mne  # Import required Python modules
**import** operator   # For the ‘add’ operator
**from** functools **import reduce**
~~~

~~~
# Connectivity objects are morphed back to the fsaverage brain
fsaverage = mne.read_source_spaces(‘fsaverage-src.fif’)
# For each of 20 subjects, read connectivity for different conditions.
# Re-order the vertices to be in the order of the fsaverage brain.
face = []
scrambled = []
contrast = []
**for** subject **in** [‘sub%03d’ % (i + 1) **for** i **in range**(20)]:
      con_face = conpy.read_connectivity(‘%s-face-con.h5’ % subject)
      con_face = con_face.to_original_src(fsaverage)
      con_scrambled = conpy.read_connectivity(‘%s-scrambled-con.h5’ % subject)
      con_scrambled = con_scrambled.to_original_src(fsaverage)
      face.append(con_face)
      scrambled.append(con_scrambled)
      contrast.append(con_face - con_scrambled) # Create contrast
# Compute the grand-average contrast
contrast = **reduce**(operator.add, contrast) / 20
# Perform a permutation test to only retain connections that are part of a
# significant bundle.
connection_indices = conpy.cluster_permutation_test(
      face, scrambled,           # The two conditions
      cluster_threshold=4,      # The initial t-value threshold to form bundles
      max_spread=0.013,          # Maximum distance (in m) between connections
                                    # that are assigned to the same bundle.
      src=fsaverage,             # The source space for distance computations
      n_permutations=1000,      # The number of permutations for estimating
                                    # the distribution of t-values.
      alpha=0.05                  # The p-value at which to reject the null-hypothesis
)
# Prune the contrast connectivity to only contain connections that are part of
# significant bundles.
contrast = contrast[connection_indices]
~~~

### Application to the example dataset

In the connectivity analysis of the example data, we focus on a selection of connections that show the most reliable difference between the experimental conditions. The pruning of the all-to-all connectivity results is implemented in script `12_connectivity_stats.py`.

In our analysis of the example dataset, we applied an initial *t*-value threshold of 5 to the connections, retaining 849 out of the total of 4781057 connections. During the clustering step, connections with start and end points within 1 cm were grouped, resulting in 144 bundles. The above thresholds were chosen such that there remained a small and manageable subset of the full all-to-all connectivity network, which shows the most robust differences between the processing of faces versus scrambled images. The permutation test revealed two bundles that show a significant difference in coherence between the processing of faces versus scrambled images (*p* < 0.05), containing a total of 240 connections.

## Visualization

Depending on the statistical threshold, there may be hundreds or thousands of connections that survive the pruning step. In order to visualize this many connections, we use a combination of a circular connectogram that summarizes connectivity between parcels (i.e., predefined cortical regions based on a brain atlas), and a “degree map” that shows, for each source point, the total number of connections from and to the point. In this framework, we may use the circular connectogram to assess global connectivity patterns between parcels and use the degree map to see which specific parts of the cortex contain the start and end points of the connections.

To create a connectivity object that defines connectivity between parcels, rather than source points, we use brain atlases, such as the ones provided by the FreeSurfer package. These atlases provide a list of parcels (also referred to as “labels”) and a list of vertices of the cortical mesh belonging to each parcel. Using this information, we can determine which source points belong to which parcel and make a parcel-wise summary.

In our pipeline, we choose to summarize the connection between two parcels by counting the total number of connections between them that survived the statistical thresholding (i.e., the degree). The summary can then be visualized using a circular connectogram. In general, large parcels that contain many source points will have more connections and thus a larger degree. Therefore, if the intention is for the circular connectogram to represent the overall connectivity between parcels, this “degree bias” could lead to misinterpretation of the result and it may be appropriate to remove this bias. This can be done by dividing the sum by the total number of possible connections from and to the parcel.

The cortical degree map is created by counting the number of connections that survived the statistical threshold from and to each source point. This degree map suffers from a similar bias as the circular connectogram, so it may be appropriate to divide the initial summary of each source point by the total number of possible connections from and to the point to remove this bias.

### Code example

The following example will parcellate a connection object according to the “aparc” brain atlas,^35^ create a circular connectogram and a cortical degree map:

~~~
**import** conpy, mne # Import required Python modules
con = conpy.read_connectivity(‘contrast-con.h5’) # Load connectivity object
l = mne.read_labels_from_annot(‘fsaverage’, ‘aparc’) # Get parcels from atlas
**del** l[-1] # Drop the last parcel (unknown-lh)
# Parcellate the connectivity object and correct for the degree bias
con_parc = con.parcellate(l, summary=‘degree’, weight_by_degree=True)
con_parc.plot() # Plot a circle diagram showing connectivity between parcels
# Plot a vertex-wise degree map and connect for the degree bias
brain = con.make_stc(‘degree’, weight_by_degree=True).plot(hemi=‘split’)
brain.add_annotation(‘aparc’) # Draw the ‘aparc’ atlas on the degree-map
~~~

The above example results in a very basic circular connectogram. For optimal clarity, some care needs to be put into the order and organization of the parcels along the circle. For example, it may be useful to dedicate the left half of the circle to parcels in the left hemisphere and the right half to the right hemisphere. Script `figure_connectivity.py` contains a more elaborate example of a circular connectogram.

### Application to the example dataset

The visualization of the pruned all-to-all connectivity of the example dataset is implemented in script `figure_connectivity.py` and presented in figure 4.

**Figure 4:**
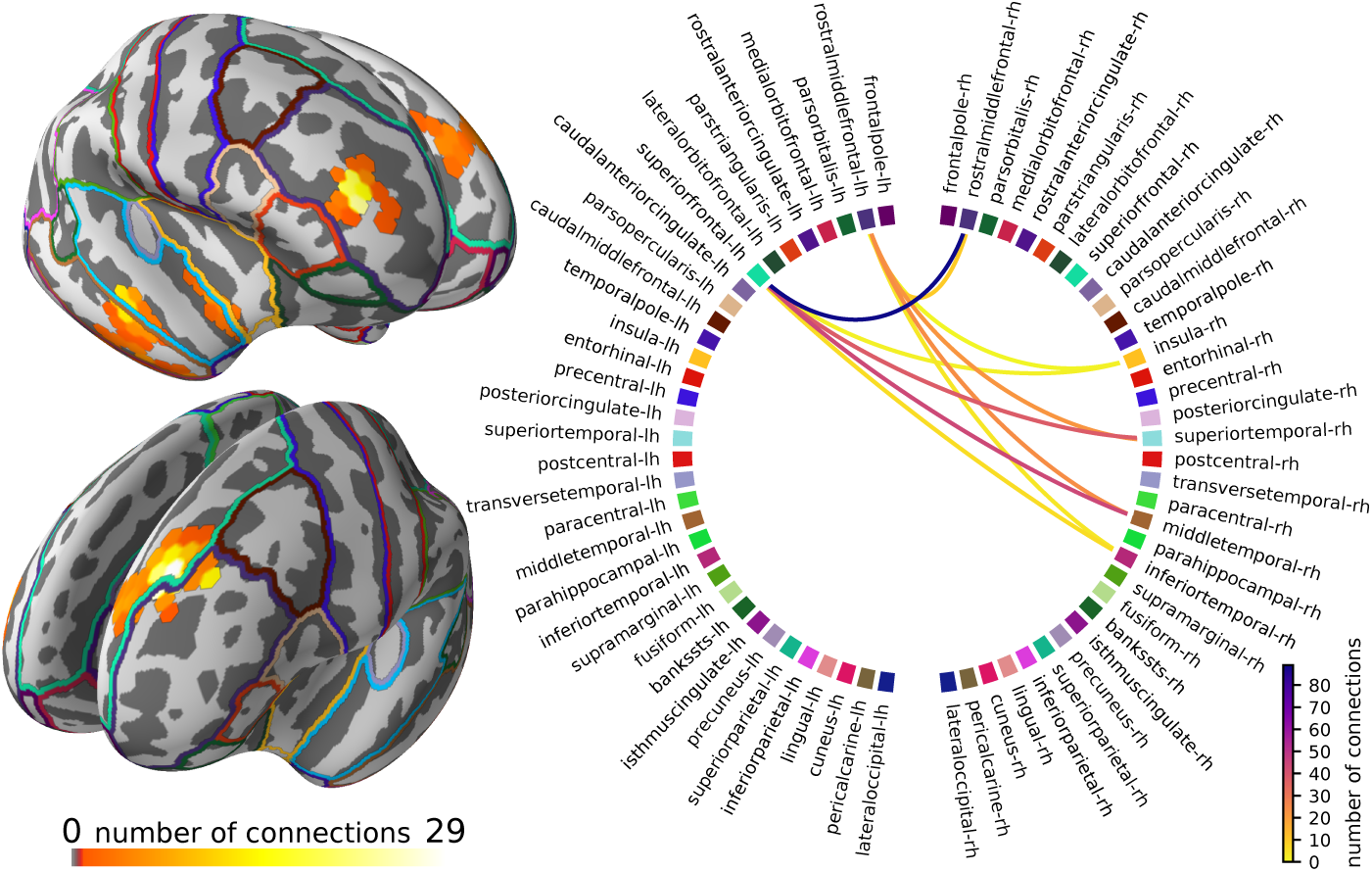
The subnetwork of the all-to-all connectivity network that shows the most robust changes across the experimental conditions. **Left:** degree map showing, for each source point, the percentage of connections, out of all possible connections, that survived the statistical threshold and clustering operations. **Right:** circular connectogram showing the number of connections between each parcel. Parcels were defined using the “aparc” anatomical brain atlas, provided by FreeSurfer.

The intended interpretation of figure 4 is to first, using the degree map, identify the main areas where connectivity changes between faces versus scrambled images and then see which parcels overlap with these areas. Then, using the circular connectogram (figure 4, right), we can determine which connections between these areas are influenced by the experimental manipulation.

In our example dataset, the pruning of the all-to-all connectivity network resulted in a subnetwork that highlights communication between the right temporal pole, right prefrontal cortex and left superior frontal cortex (figure 4, top-left). Since the obtained connectome is so sparse, we opted not to compensate for the degree bias in the degree map and circular connectogram and simply report the number of connections.

The start and end points of the connection bundles do not always line up well with the parcels that are defined by the “aparc” brain atlas, which makes it less obvious in the circular connectogram that we are looking at two bundles of connections. However, when the circular connectogram is interpreted alongside the degree map, the two bundles become clear.

## Discussion

The presented analysis pipeline facilitates mapping of cortico-cortical coherence, specifically its modulation between experimental conditions, in an all-to-all manner based on whole-head MEG data. The original estimation of coupling is conducted at the level of a detailed grid of source points covering the entire cortex, but statistical testing and visualization of the results can be conducted both at this level and at the level of a coarser cortical parcels. In addition to the estimation of connectivity, the pipeline provides source estimates of oscillatory activity (“power mapping”) at the same spatial scales as used in the coherence analysis. The analysis pipeline consists of several steps that involve choices regarding how connectivity can be estimated, some of which are general considerations that are relevant also for other pipelines than the one presented here. In this section, we discuss the effects and possible developments regarding some of these choices for the most critical analysis steps.

### Estimation of the cross spectral density matrix

In the present manuscript, we considered cortico-cortical connectivity for event-related experimental paradigms where the cross spectral density matrix, which represents the mutual dependencies of neural signals at the sensor-level, needs to be estimated in a time-resolved manner. This type of analysis is useful as it allows the use of event-related paradigms where experimental manipulation is generally more straightforward than in continuous and more naturalistic experiments. Moreover, the approach readily allows limiting the analysis to an artefact-free time window of the experiment (e.g., in speech production). The original DICS was developed for continuous data^36^ where the CSD estimation is based on Fourier transformations. In the present analysis, as well as in previous work using event-related DICS,^37^ wavelet-based analysis was used to obtain the time-frequency CSD. In the time-frequency domain, wavelets provide an optimal compromise between time and frequency resolution. However, the time-resolved estimation could equally well be conducted using short-term Fourier transformation, especially if appropriate windowing functions are used. More importantly, while the present analysis focused on the event-related estimation of cortico-cortical coupling, the pipeline can directly be applied also to continuous data by replacing the CSD estimation step with Fourier transformation based computations, as was done in Gross et al. (2001).

### Definition of the source space

In general, reliable evaluation of cortico-cortical connectivity requires a group-level description of neural interactions. This, in turn, necessitates the estimation of the neural connectivity patterns in the same locations across subjects. This can be achieved both at the level of detailed grids of source points and cortical parcellations. Parcellations have been used more commonly for all-to-all type connectivity estimation^38^ as they reduce the computational load of the estimation and the amount of statistical testing. The present analysis pipeline facilitates both using a grid of source points and parcel-level estimation. An effective group-level estimation of connectivity between source points is achieved by generating a grid of points along the cortex of a reference brain (e.g. FreeSurfer’s “fsaverage” brain) and transforming this grid to each individual’s anatomy. As a consequence, the same connections are estimated in every subject, allowing direct estimation of the group-level statistics. A parcel-level description can then also by readily obtained as it is sufficient to assign each point-level connection to a parcel-pair in the common brain instead of doing the assignments separately in each subject. For the parcel-level estimation it would be almost equally straightforward to use individually defined grids of source points. However, when one aims to evaluate more detailed spatial aspects of connectivity, the chosen approach eliminates the need for massive interpolation operations that would be required if individual-level grids of source points were used.

### Choice of the interaction metric

Here, we chose to apply a DICS based estimation of cortical connectivity that allows a direct mapping of the mutual dependencies of the sensor-level signals to a cortical space without the need for estimation of cortical-level time-series of activity. As the present connectivity estimation is dependent on the use of a CSD matrix, coherence is the only interaction metric that can be estimated straightforwardly in this manner. Notably, similar approaches that map the sensor level interaction patterns to the source level without the time-series estimation step have also been developed for metrics such as partial directed coherence^39^ or imaginary coherence.^40^

Since interactions due to field spread exhibit zero phase lag, using an interaction measure that is sensitive only to non-zero phase lag, such as imaginary coherence,^41^ may reduce the detection of spurious interactions. However, there is good indication that not all zero-phase-lag connections are spurious,^42^ so methods focusing solely on imaginary coherence should be used with care.

As theoretical models of neural interactions propose that neuronal coherence mechanistically subserves neuronal communication,^43^ the choice of coherence as an interaction metric factor does not necessarily represent a limitation of the approach. However, if the goal is to use some other metric to quantify neural interactions the analysis pipeline would need to be adjusted. Within the framework of transforming sensor-level dependency patterns to the source level, it would be possible to utilize, e.g., weighted phase-lag index^44^ by transforming single-trial (as opposed to average) CSD matrices to the source level. Most metrics would, however, require that one would first estimate cortical-level time-series of activity. This would be readily possible by using the DICS spatial filter for weighing the sensor-level time-series. Within this framework, however, the use of a detailed grid of source points would no longer be computationally tractable and it would be better to construct parcel-level time-series before the metric-dependent quantification of neural interactions.

### Considerations regarding field spread and source orientations

In the presented analysis we focused on estimating connectivity in an all-to-all manner without the need for a priori seed regions or constraining of the analysis to connections between preselected brain areas. The field spread based confounding factors in connectivity estimation are particularly critical for this type of analysis^45^ where it is difficult, e.g., to visually evaluate whether the observed changes in patterns of neural interactions truly represent modulation of coupling as opposed to modulation of field spread between experimental conditions. To minimize the effects of field spread, we focused only on long range (≥4 cm) connections and examined coherence modulations for conditions for which the amount of neural activity/oscillatory power are closely matched. However, the exact degree of matching that is required for contrasting canonical coherence, or any other type of coherence estimates, remains an open question.

Most of the neural signals detected by MEG and EEG originate from sources that are approximately orthogonal to the surface of the cortex.^46^ So, one may restrict the source space by defining only ECDs that point in the orthogonal direction, by for example leveraging the surface normals of the 3D-mesh produced by FreeSurfer.^47^ However, in practice, each source grid-point represents the signal for a patch of cortical surface which, due to the folding of the cortex, includes locations with different surface normals. Especially when using a large spacing between grid-points, as we do in our pipeline, the source within each patch that drives the activity at the grid-point does not necessarily have the same orientation as the average surface normal of the patch as a whole. This is why it is recommended to allow for some flexibility regarding dipole orientation whenever possible.^48^ Whether this is possible in practice depends on the computational costs of performing the source estimates for multiple orientations and whether the SNR is good enough to produce a reliable estimate of the optimal orientation.

In the current pipeline we exclusively used a canonical estimation of coherence^49^ where, for each connection, the orientations of the source ECDs at both sides of the connection are selected such that they maximize coherence. To make this computationally feasible, we restrict the number of possible orientations by leveraging the fact that MEG is less sensitive to “radial” sources, due to the properties of the magnetic field.^50^ By choosing a tangential source space (see figure 2, right), coherence values are only computed for those ECD orientations that yield the largest signal on the MEG sensors. This canonical estimation of coherence yields a maximally stable estimate of coherence and it is well suited for investigating modulation of coherence between experimental conditions. The estimates are, however, relatively smooth. For estimating absolute coherence values for short-range connections, especially when the expected coherence values are small, other criteria for defining the source orientations could be more appropriate.

For the connectivity analysis, we chose to design separate sets of DICS beamformer filters for each condition, instead of designing one set of filters to apply to both conditions. Accordingly, the estimation of coherence and optimization of the source orientations was also performed separately for each condition. This approach allows for subtle differences in optimal source orientations between the conditions and avoids biasing the solution towards the condition with better SNR. If the goal were to ensure that field spread effects are be maximally cancelled out by contrasting two conditions, it would be beneficial to conduct the orientation optimization and weight estimation using a joint CSD across the conditions. The optimal choice between the alternatives depends on the research question and properties of the data.

### Statistical testing and visualization

In the final stage of connectivity analysis one also needs to consider both what type of statistical testing and what spatial scales are optimal. As stated above, the present analysis pipeline has been designed for examining coherence modulations between experimental conditions. Moreover, to minimize confounding effects resulting from substantial power differences between conditions, the pipeline is aimed at contrasting different tasks as opposed to contrasting a single task to resting baseline levels of neural interaction. It is also possible to contrast a single task to a task-average^51^ to highlight how the connectivity changes in a specific task with respect to multiple different tasks. Notably, the analysis pipeline does not provide a full connectome, that is, a complete description of the underlying networks. Instead, it yields a snapshot of a specific part of the network where cortico-cortical coupling has changed from one experimental condition to another. By introducing a battery of control conditions and comparisons between different conditions, the pipeline would thus allow the identification of different subnetworks that are critical for different aspects of neural processing in performing the tasks.

The present analysis pipeline enables the evaluation of the above aspects both at the level of detailed grids of source points and coarser parcellations. An effective visualization combines a connectogram that shows the connectivity at the parcel level, with a visualization of the grid-level connectivity on the cortex. This makes it possible to evaluate whether the patterns of neural connectivity evaluated at the grid-level are faithfully represented also at the level of a parcellated cortex. This, in turn, allows the fine tuning of the parcellation schemes, which are generally based on anatomical division, to better suit MEG data.

## Conclusion

We have presented an analysis pipeline that facilitates the cortical mapping of oscillatory activity and estimation of all-to-all type cortico-cortical coherence. Combined with Jas et al. (2017), all the necessary steps of the analysis of a real experiment are described: starting from the processing of raw MEG data to the statistical group analysis of the networks and visualization of the results using connectograms, as one would use in a publication. We have developed a new python package called ConPy, which integrates with MNE-python^52^ to offer a clean interface to all required software routines to reproduce the analysis. It is our hope that our example analysis will serve as a strong foundation for others who seek to implement their own DICS analysis pipelines.

## Author Contributions

MvV, ML, RS, JK: writing of the manuscript. ML, RS, JK: conceptualization of the DICS analysis pipeline. MvV, SA: software implementation of the analysis pipeline. MvV: analysis of the example dataset.

## Acknowledgments

We would like to thank Britta Westner, Sarang S. Dalal, Eric Larson and Alexandre Gramfort for their help with reviewing the code.

## Footnotes

1 Van Veen, van Drongelen, Yuchtman, and Suzuki, 1997

2 Gross et al., 2001; Jan Kujala, Gross, and Salmelin, 2008

3 Liljeström, Kujala, Stevenson, and Salmelin, 2015; Betti et al., 2013; Gonzalez-Castillo and Bandettini, 2017; Liljeström, Vartiainen, Kujala, and Salmelin, 2018

4 Hannu Laaksonen, Kujala, Hultén, Liljeström, and Salmelin, 2012; Salmelin and Kujala, 2006

5 J. M. Schoffelen and Gross, 2009

6 Gross et al., 2001

7 Bressler and Kelso, 2001; Fries, 2005

8 Buffalo, Fries, Landman, Buschman, and Desimone, 2011; Donner and Siegel, 2011; Hipp, Hawellek, Corbetta, Siegel, and Engel, 2012; Liljeström, Kujala, et al., 2015

9 Jan Kujala et al., 2008

10 Liljeström, Kujala, et al., 2015; Saarinen, Jalava, Kujala, Stevenson, and Salmelin, 2015

11 Hannu Laaksonen, 2012

12 Liljeström, Kujala, et al., 2015; Liljeström, Stevenson, Kujala, and Salmelin, 2015

13 Gramfort et al., 2013

14 Saarinen et al., 2015

15 J. M. Schoffelen and Gross, 2009

16 Wakeman and Henson, 2015

17 Wakeman and Henson, 2015

18 H. Laaksonen, Kujala, and Salmelin, 2008

19 Saarinen et al., 2015; Alexandrou, Saarinen, Mäkelä, Kujala, and Salmelin, 2017

20 Dale, Fischl, and Sereno, 1999

21 Dale et al., 1999

22 Fischl, Sereno, Tootell, and Dale, 1999

23 Fischl et al., 1999

24 Hämäläinen, Hari, Ilmoniemi, Knuutila, and Lounasmaa, 1993

25 Hämäläinen et al., 1993

26 Van Veen et al., 1997

27 Jas et al., 2017

28 Ciulla, Takeda, and Endo, 1999

29 Fries, 2005

30 Hämäläinen et al., 1993

31 Salmelin and Kujala, 2006

32 Gross et al., 2001

33 Rosazza and Minati, 2011

34 Maris and Oostenveld, 2007

35 Fischl et al., 2004

36 Gross et al., 2001

37 Jan Kujala, Vartiainen, Laaksonen, and Salmelin, 2012; J. Kujala et al., 2014; Liljeström, Kujala, et al., 2015

38 Palva, Monto, Kulashekhar, and Palva, 2010; Saarinen et al., 2015; J.-M. Schoffelen et al., 2017

39 Michalareas, Schoffelen, Paterson, and Gross, 2013

40 Drakesmith, El-Deredy, and Welbourne, 2013

41 Nolte et al., 2004; Drakesmith et al., 2013

42 Gollo, Mirasso, Sporns, and Breakspear, 2014

43 Fries, 2005

44 Vinck, Oostenveld, van Wingerden, Battaglia, and Pennartz, 2011

45 J. M. Schoffelen and Gross, 2009

46 Hämäläinen et al., 1993

47 Dale and Sereno, 1993

48 Lin, Belliveau, Dale, and Hämäläinen, 2006

49 Saarinen et al., 2015; Liljeström, Kujala, et al., 2015; Liljeström, Stevenson, et al., 2015

50 Hämäläinen et al., 1993

51 Saarinen et al., 2015

52 Gramfort et al., 2013

